# Midkine-a regulates the formation of a fibrotic scar during zebrafish heart regeneration

**DOI:** 10.1101/2021.01.29.428781

**Authors:** Dimitrios Grivas, Álvaro González-Rajal, José Luis de la Pompa

**Affiliations:** Intercellular Signalling in Cardiovascular Development and Disease Laboratory, Centro Nacional de Investigaciones Cardiovasculares (CNIC), Melchor Fernández Almagro 3, Madrid 28029, SPAIN; Ciber de Enfermedades Cardiovasculares, 28029 Madrid, SPAIN; Cell Division Lab, ANZAC Research Institute, Gate 3, Hospital Road, Concord 2139, NSW, AUSTRALIA

## Abstract

The adult zebrafish heart regenerates after injury, unlike the hearts of mammals. Heart cryoinjury triggers the formation of a fibrotic scar that gradually degrades, leading to regeneration. Midkine-a (Mdka) is a multifunctional cytokine that is activated after cardiac injury. Here, we investigated the role of *mdka* in zebrafish heart regeneration. We show that *mdka* expression is strongly induced at 1-day post cryoinjury (dpci) throughout the epicardium, whereas by 7 dpci expression has become restricted to epicardial cells covering the injured area. To study the role of *mdka* in heart regeneration, we generated *mdka*-knock out (KO) zebrafish strains. Analysis of injured hearts showed that loss of *mdka* decreased endothelial cell proliferation and resulted in a blockade of heart regeneration characterized by retention of a collagenous scar. Transcriptional analysis revealed increases in collagen transcription and TGFβ signalling activity. These results reveal a critical role for *mdka* in fibrosis regulation during heart regeneration.

## Introduction

The adult mammalian heart has limited regeneration capacity, and myocardial infarction (MI) generates a permanent fibrotic scar that progressively leads to heart failure (Benjamin et al., 2019). Conversely, zebrafish can regenerate damaged cardiac tissue (Poss, 2002). Heart cryoinjury results in massive cell death and the formation of scar tissue that gradually resolves, resulting in complete heart regeneration within 90 days (Chablais and Jaźwińska, 2012; González-Rosa and Mercader, 2012; Schnabel et al., 2011). The most abundant extracellular matrix (ECM) component responsible for fibrotic tissue formation is collagen (Theocharis et al., 2016). Fibrosis progression is dependent on the ECM molecule periostin, which promotes collagen cross-linking and fibroblast activation (Ito et al., 2014; Sánchez-Iranzo et al., 2018). In the injured zebrafish heart, the main source of ECM molecules are epicardial cells and epicardial-derived fibroblasts (Marro et al., 2016; Sánchez-Iranzo et al., 2018; Wang et al., 2013). TGFβ signalling, important for matricellular protein production, is active during regeneration, and chemical inhibition of the pathway decreases collagen deposition and blocks regeneration (Chablais and Jazwinska, 2012). Heart regeneration is also prevented by inhibition of fibrotic tissue resolution (Xu et al., 2018), indicating that heart regeneration requires both formation and removal of scar tissue.

*Midkine* (*mdk*) is a small, pleiotropic cytokine that is upregulated after injury in several tissues, including the heart and retina (Lien et al., 2006; Luo et al., 2012). Cardiac insult in *Midkine-KO* mice leads to larger injuries and decreased heart function (Horiba et al., 2006; Ishiguro et al., 2011). Prolonged treatment of Midkine to injured hearts results in reduced collagen deposition in both the infarcted and healthy areas of the ventricle (Fukui et al., 2008; Sumida et al., 2010; Takenaka et al., 2009). Additionally, Midkine improves angiogenesis in injured cardiac tissue (Fukui et al., 2008; Sumida et al., 2010; Takenaka et al., 2009), indicating the importance of Midkine in the cardiac response to injury in mammals. In zebrafish, *mdka* is upregulated after injury in several tissues, including the heart, fin, and retina. In the absence of *mdka*, Muller glia proliferation is decreased leading to impaired photoreceptor regeneration after retina injury (Nagashima et al., 2020). Additionally, fin regeneration is delayed due to an initial decrease in cell proliferation, although ultimately the fin is fully regenerated (Ang et al., 2020). Despite its importance during the cardiac response to injury in mammals, little is known about the role of *mdka* in zebrafish heart regeneration.

Here, we generated *mdka-KO* zebrafish and studied heart regeneration. We found that *mdka* is upregulated upon injury in the epicardium. Deletion of *mdka* decreased endothelial cell proliferation and prevented heart regeneration, leading to an injured area enriched in collagen deposition. Transcriptional analysis revealed upregulation of TGFβ signalling activity and expression of ECM molecules, including collagen and periostin, revealing a crucial role for *mdka* in the regulation of the fibrotic response upon heart injury.

## Results and Discussion

### *mdka* expression in the regenerating heart

The expression of *mdka* is upregulated upon heart injury in several animal models (Fukui et al., 2008; Horiba et al., 2006; Lien et al., 2006; Sumida et al., 2010). We performed a quantitative PCR (qPCR) analysis in cryoinjured zebrafish hearts to analyse *mdka* expression during regeneration (Figure 1A). *mdka* RNA was strongly expressed in the heart at 1 dpci, reaching a peak at three days post cryoinjury (dpci), followed by a reduction at 7 dpci. We next investigated the expression pattern of *mdka* by *in situ* hybridization (ISH) (Figure 1B-D’). At 1 dpci, *mdka* expression was detected in the epicardium, with no expression in intact hearts (Figure 1_Supplementary Figure 1 A, A’, Figure 1B, B’). At 3 dpci, *mdka* expression was upregulated in the epicardium, and by 7 dpci the expression became restricted to the multi-layered epicardium covering the injury site (Figure 1C-D’). Epicardial cells maintained *mdka* expression at 14 dpci (Figure 1_Supplementary Figure 1B, B’), and *mdka* remained detectable up to 130 dpci in the epicardium, the area between the cortical and trabecular myocardium, and in the compact layer, hinting at vascular expression (Figure 1_Supplementary Figure 1C, C’). To further define the epicardial localization of *mdka*, we performed fluorescence ISH in heart sections of *Tg(wt1b:GFP)* zebrafish, which express GFP in *wt1b*^+^ epicardial cells after injury (Gonzalez-Rosa et al., 2011). *mdka* ISH was followed by immunostaining for GFP and Aldh1a2 (Kikuchi et al., 2011a) (Figure 1E-F’’). *mdka* was expressed in GFP^+^ cells (Figure 1F, F’’, arrowheads) and in Aldh1a2 expressing cells (Figure 1F’, F’’, arrows), indicating that injury triggers *mdka* expression in signalling cells. Our analysis shows that *mdka* is upregulated in the epicardium upon injury and later persists in the activated epicardium. By 7 dpci, the epicardium comprises a heterogeneous cell population that includes macrophages and fibroblasts (Sánchez-Iranzo et al., 2018; Sanz-Morejón et al., 2019). The main source of ECM is fibroblasts (Kendall and Feghali-Bostwick, 2014), and RNA-seq data from *postnb*^*+*^ cells shows that *mdka* is highly upregulated in epicardial-derived fibroblasts in the regenerating zebrafish heart (Sánchez-Iranzo et al., 2018). *mdka* is also widely expressed in the epicardium, that secretes ECM components (Wang et al., 2013), suggesting that *mdka* is involved in the fibrotic response after injury.

**Figure 1.**
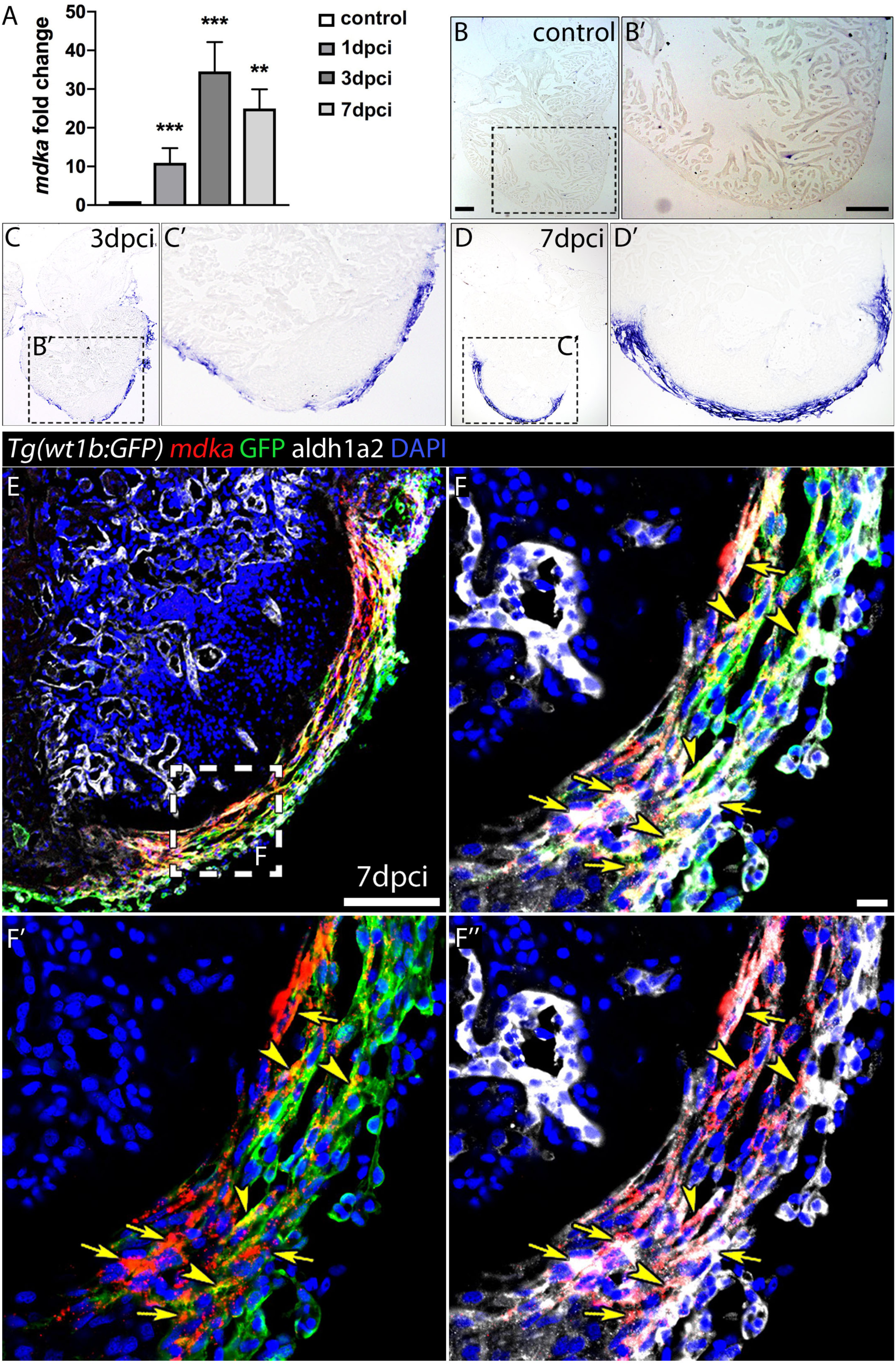
*mdka* expression upon injury. (A) qPCR analysis of *mdka* in regenerating hearts. ANOVA tests; \*\**P*<0.01, \*\*\**P*<0.001; Mean±S.D. (B-D’) ISH of *mdka* in intact (B, B’), 3 dpci (C, C’), and 7 dpci (D, D’) hearts. Scale bars, 100 µm. (E-F’’) Fluorescent ISH for *mdka* followed by immunolabelling for GFP and Aldh1a2 in 7 dpci *Tg(wt1b:GFP)* hearts. Arrowheads indicate *mdka*-GFP overlap and arrows *mdka*-Aldh1a2 overlap. Scale bar E, 100 µm; F’-F’’, 10 µm.

### *mdka-KO* hearts fail to regenerate

Our initial analysis showed that *mdka* expression is readily activated in the epicardium after heart injury (Figure 1). Since *mdka* is not expressed in uninjured adult hearts but is highly upregulated upon injury, we reasoned that loss of *mdka* may have a significant effect on heart regeneration. Therefore, we generated *mdka* knock-out (KO) zebrafish (Figure 2 and Figure 2_Supplementary Figure 1). The mutant allele *mdka*^*cn105*^ decreased significantly *mdka* expression without upregulation of *mdka* paralogue, *mdkb* (Figure 2A) and resulted in complete loss of Mdka protein as revealed by immunofluorescence and western blot analysis (Figure 2B-D). We next examined the effect of *mdka* inactivation on heart regeneration. Hearts from *mdka*^*+/+*^, *mdka*^*cn10*5^, and *mdka*^*+/cn105*^ animals were cryoinjured, and analysed after 90-days (Figure 2E-G). The analysis revealed that *mdka*^*cn105*^ hearts had significantly larger injuries than their *mdka*^*+/+*^ and *mdka*^*cn105/+*^ counterparts (Figure 2H). Additionally, the injured area of *mdka*^*cn105*^ hearts was characterized by persistent collagen deposition compared with *mdka*^*+/+*^ and *mdka*^*cn105/+*^ hearts (Figure 2I). These results indicate that *mdka* deletion blocks heart regeneration.

**Figure 2.**
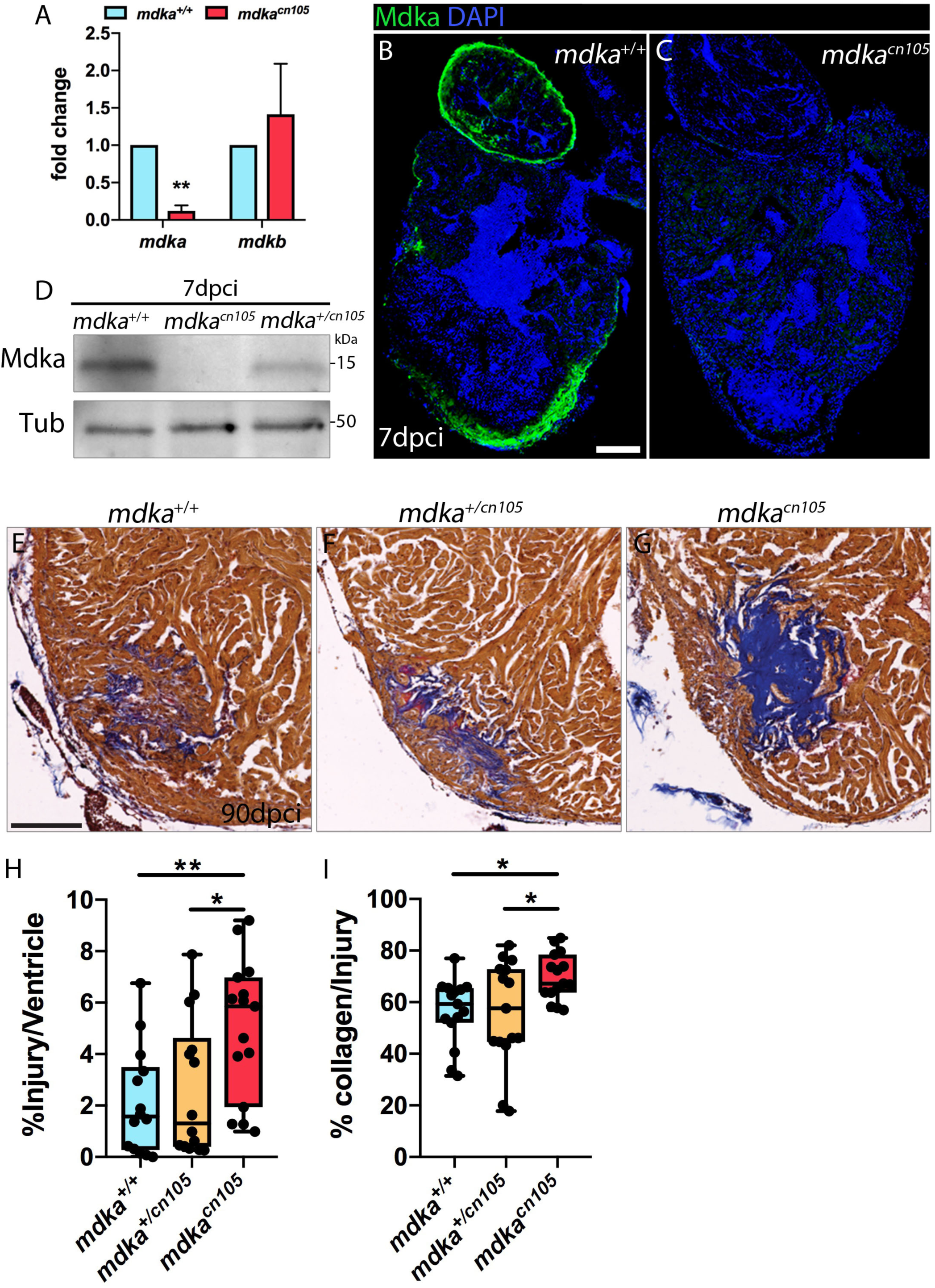
Loss of *mdka* leads to arrest of heart regeneration. (A) qPCR analysis of *mdka* and *mdkb* in 2 dpf *mdka*^*+/+*^ and *mdka*^*cn105*^ embryos. t-test; \*\**P*<0.01; Mean±S.D. (B, C) Immunofluorescence staining of Mdka in 7 dpci *mdka*^*+/+*^ and *mdka*^*cn105*^ hearts. Scale bar, 100 µm. (D) Western blot (WB) analysis for Mdka in 7 dpci *mdka*^*+/+*^, *mdka*^*cn105*^, and *mdka*^*+/cn105*^ hearts. Tub, a-Tubulin; kDa, kilodalton. (E-G) AFOG staining of 90 dpci *mdka*^*+/+*^, *mdka*^*+/cn105*^, and *mdka*^*cn105*^ hearts. Collagen in blue, fibrin in red, healthy myocardium in brown. Scale bar, 100 µm. (H) Quantification of injury size calculated as damaged tissue (collagen and fibrin) as a percentage of the ventricular area. n_WT_ = 14, n_KO_= 15, n_HET_= 14. (I) Collagen as a percentage of the injured area. n_WT_ = n_KO_= n_HET_= 15. ANOVA; \**P*<0.05; \*\**P*<0.01; Mean±S.D.

### Increased fibrosis in injured *mdka*^*cn105*^ hearts

To further study the effect of *mdka* deletion, we compared the transcriptional profile of 7 dpci *mdka*^*+/+*^ and *mdka*^*cn105*^ ventricles by RNA-seq (Figure 3A, Supplementary Table 1). Ingenuity pathway analysis (IPA) identified enrichment of processes involved in heart failure, including cardiac fibrosis, heart degeneration, and vascular lesions (Figure 3B). Analysis of upstream regulators revealed increased activation of the TGFβ pathway, including Tgfβ1, Tgfbr1, and Smad downstream effectors such as Smad3 and Smad4, whereas the TGFβ inhibitory effector Smad7 was reduced (Figure 3C) (Moustakas and Heldin, 2009).

**Figure 3.**
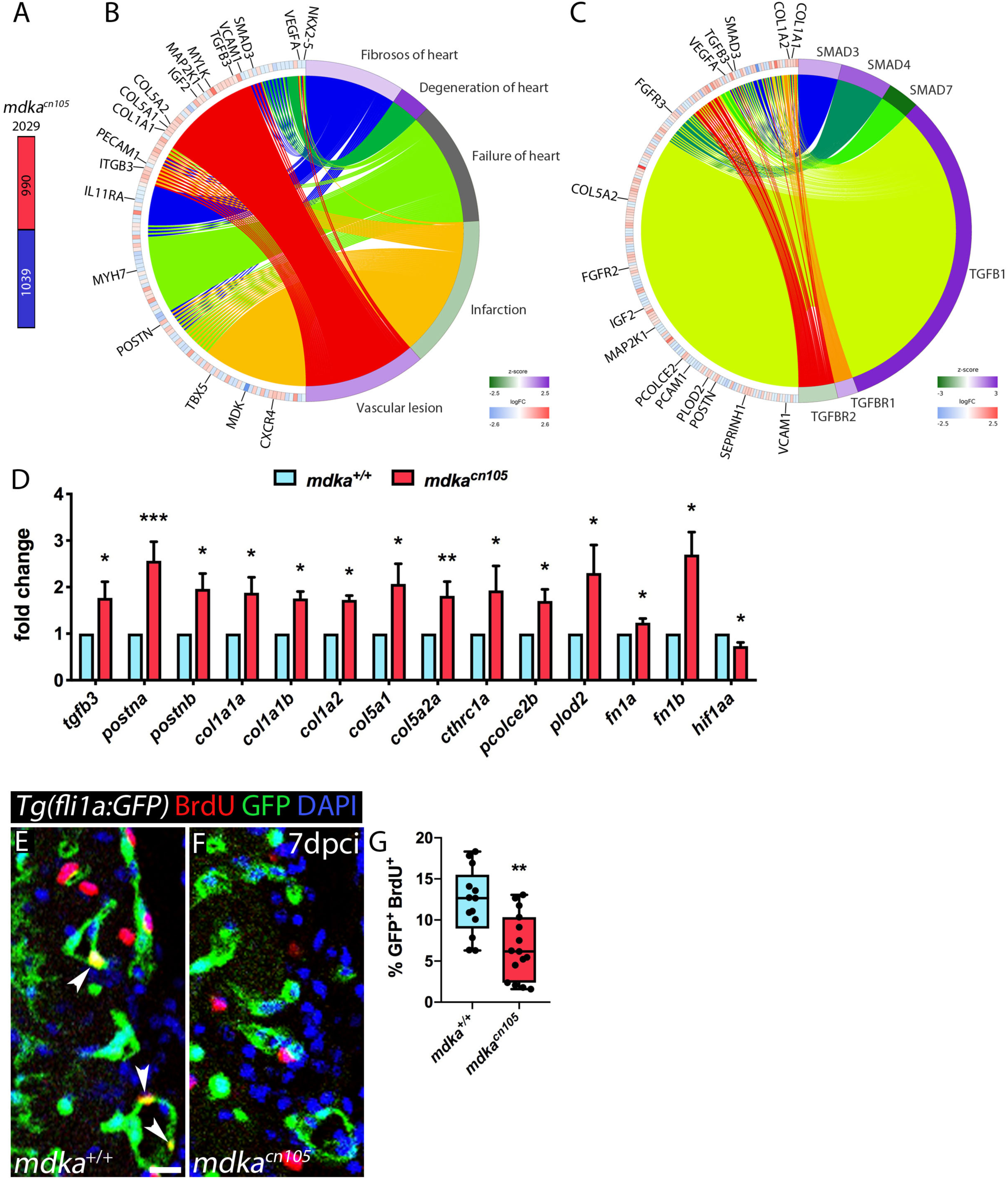
Transcriptional profiling of regenerating *mdka*^*cn105*^ hearts. (A) Total number of differentially expressed genes identified by RNA-seq. Numbers indicate upregulated genes (red) and downregulated genes (blue). (B, C) Circular plots showing representative differentially expressed genes (left semicircle perimeter) and IPA biofunctions and upstream regulators (right semicircle perimeter). Activation z-score scale: green, repression; magenta, activation; white, unchanged. LogFC scale: red, upregulated; blue, downregulated; white, unchanged. (D) qPCR analysis of *mdka*^*+/+*^ and *mdka*^*cn105*^ 7 dpci hearts. t-test; *\*P*<0.05,\*\**P*<0.01, *\*\*\*P*<0.001; Mean±S.D. (E, F) Immunofluorescence staining of GFP and BrdU in 7 dpci *Tg(fli1a:GFP)* heart sections. Scale bar, 10 µm. (G) Quantification of BrdU^+^/GFP^+^ endothelial cells. n_WT_ = 13, n_KO_ = 15; t-test; *\*\*P*<0.01; Mean±S.D.

In *mdka*^*cn105*^ hearts, genes associated with fibrosis were significantly upregulated (Figure 3_Supplementary Figure 1). These included genes for the ECM components *col1a1a, col1a1b, col1a2, col5a1, col5a2a*, and *postna* (Chablais and Jazwinska, 2012; Ito et al., 2014). Also upregulated were *pcolce2b, cthrc1a* and *plod2*, which are involved in collagen stability and scar tissue formation. *pcolce2b* is involved in collagen-fibril assembly, ECM formation and *Tgfβ1* stimulation in pancreatic fibrosis, and is expressed in the epicardium after heart injury in zebrafish (Cao et al., 2016; Sorci-Thomas et al., 2015). *cthrc1a* contributes to fibrosis progression via TGFβ-dependent enhancement of liver fibrosis (Li et al., 2019), and *plod2* is involved in collagen fiber architecture and fibrosis development (Gilkes et al., 2013). In contrast, expression of *vegfaa* and *hif1aa*, genes required for vascular development and repair, was decreased in *mdka*^*cn105*^ hearts (Gerri et al., 2017; Marín-Juez et al., 2016; Marín-Juez et al., 2019).

We validated our RNA-seq results by qPCR analysis that further showed increased expression of *postnb, fibronectin1a*, and *fibronectin1b* (Figure 3D). We confirmed the transcriptional upregulation of several collagens, including the fibrillar collagens *col1a1a, col1a1b, col1a2, col5a1*, and *col5a2a*, which associate with type I fibrils (Figure 3D) (Theocharis et al., 2016). Upon heart cryoinjury in zebrafish, *tgfβ1* is expressed in the injured area, and inhibition of TGFβ signalling leads to decreased collagen deposition and impairs regeneration (Chablais and Jazwinska, 2012). Collagen is the main ECM scaffolding protein, and our analysis shows that collagen is upregulated in injured *mdka*^*cn105*^ hearts. Specifically, fibril-forming *col1a1a* and *col1a1b* combine with *col1a2* to produce strong cross-linked fibres that contribute to scarring (Holmes et al., 2018), and these molecules are upregulated after *mdka* deletion. The matricellular molecules *postna* and *postnb* were also upregulated in *mdka*^*cn105*^ hearts (Figure 3D). Periostin is expressed by cardiac fibroblasts and contributes to fibroblast stimulation and persistency in injured hearts (Dixon et al., 2019; Sánchez-Iranzo et al., 2018). Additionally, it is involved in fibrillar collagen formation promoting fibrosis (Shimazaki et al., 2008).

Secretion of ECM depends on TGFβ, therefore, we evaluated TGFβ pathway activation by examining the downstream effector phospho-Smad3 (psmad3) (Figure 3_Supplementary Figure 2). The analysis revealed that phosphorylation of Smad3 was similar between *mdka*^+/+^ and *mdka*^*cn105*^ hearts, suggesting that *mdka* affects TGFβ activation thought non-canonical pathways. In zebrafish, formation and degradation of the scar tissue are crucial for heart regeneration (Chablais and Jazwinska, 2012; Sánchez-Iranzo et al., 2018; Xu et al., 2018). We found that *mdka* expression overlaps with the epicardial markers Wt1b and Aldh1a2 (Figure 1E-F’’), which are also co-expressed with *tcf21*; moreover, *tcf21*^*+*^ epicardial cells participate in collagen deposition (Kikuchi et al., 2011b; Sánchez-Iranzo et al., 2018; Wang et al., 2013). Moreover, transcriptome analysis of *post1b*^*+*^ fibroblasts revealed that *mdka* is highly expressed by these cells (Sánchez-Iranzo et al., 2018). This suggests that heart regeneration arrest in *mdka*-deficient zebrafish is due to an increased fibrotic response, as evidenced by upregulation of collagens and periostin. This is in consistent with findings in the injured mammalian heart, where treatment with Midkine after MI leads to decreased collagen deposition (Sumida et al., 2010; Takenaka et al., 2009). Additionally, *postnb*^*+*^ fibroblasts are detectable until 90 dpci, which could explain the persistent *mdka* expression that we observed after regeneration was complete. These data suggest that continuous *mdka* expression by epicardial cells and epicardial-derived fibroblasts throughout the course of regeneration is required for ECM regulation, and that *mdka* loss leads to increased ECM deposition, resulting in impaired regeneration.

We then examined the proliferation of epicardial cells and cardiomyocytes, but we did not detect any difference (Figure 3_Supplementary Figure 2). Since Midkine is known to act as a pro-angiogenic factor (Sumi et al., 2002), we also analysed the proliferation of cortical endothelial cells. We found that endothelial cells adjacent to the injury site proliferate less in *mdka*^*cn105*^ hearts than in control hearts (Figure 3E-G). In addition, we detected reduced transcriptional expression in *mdka*^*cn105*^ hearts of the angiogenesis promoter *hif1aa* (Figure 3D). HIF-1a positively regulate *Midkine* expression and leads to enhanced angiogenesis (Reynolds et al., 2004), and that *hif1a* zebrafish mutant present reduced endothelial cell proliferation in regenerating hearts (Marín-Juez et al., 2019). Consequently, decreased expression of *hif1aa* in *mdka*^*cn105*^ regenerating hearts explains the decline in endothelial cells proliferation. Fast revascularization provides a vascular network necessary for repopulation of the injured area. And similar to previous reports, our data show that defects in revascularization upon injury results in arrest of regeneration and maintenance of the fibrotic tissue (Harrison et al., 2015; Marín-Juez et al., 2016),

Our analysis reveals that *mdka*, readily activated upon injury, is required for zebrafish heart regeneration. We showed that loss of *mdka* leads to dysregulation of the fibrotic response and retention of the scar as well as a decrease in angiogenesis, leading to arrest of regeneration. Hence, *mdka* promotes cardiac remodelling through the regulation of collagen turnover and vascularization, highlighting the importance that regulated attenuation of ECM production is an essential step for fibrosis regression and complete regeneration.

## Material and Methods

### Zebrafish husbandry and transgenic lines

Animal studies were approved by the CNIC Animal Experimentation Ethics Committee and by the Community of Madrid (Ref. PROEX 83.8/20). Animal procedures conformed to EU Directive 2010/63EU and Recommendation 2007/526/EC regarding the protection of animals used for experimental and scientific purposes, enforced in Spanish law under Real Decreto 53/2013. Zebrafish were raised under standard conditions at 28° C as described (Kimmel et al., 1995).

### Generation of *mdka* mutant alleles

*mdka-KO* zebrafish were generated using the oligonucleotides CACCATAGAGC CACTCCGCACAGT and AAACACTGTGCGGAGTGGCTCTAT. The oligos were inserted into the pX330 vector (Cong et al., 2013), which was linearized with BbsI (New England Biolabs, Ipswich, MA). The guide mRNA was amplified with the forward primer ACGGGGTAATACGACTCACTATAGGGATAGAGCCACTCCGCACAG, which includes T7 polymerase promoter-specific sequences, and the reverse primer AAAAAGCACCGACTCGGTGCCA. The guide RNA was injected into one-cell stage embryos together with Cas9 protein (N.E. Biolabs). Mutant animals were identified by PCR using the following primers: *mdka*-Fwd TGTTATGTATGATTCTGCGAT and *mdka*-Rvs ACAGAGGCACAAAACTACCAA.

### Heart cryoinjury and fin amputation

Fish were anaesthetised by immersion in fish water containing 0.04% tricaine (Sigma-Aldrich, St Louis, MO) and placed on a wet sponge under a dissecting microscope with the ventral side exposed. The cardiac cavity was dissected using microscissors and microforceps, and the pericardium was removed. The ventricle of the heart was exposed and dried and was then touched with a copper probe previously immersed in liquid nitrogen (González-Rosa and Mercader, 2012). The fish were immediately returned to water to recover. Amputation of the caudal fin for regeneration experiments was performed as described (Munch et al., 2013). Briefly, fish were anaesthetized in 0.04% tricaine, and half of the caudal fin was amputated.

### Bromodeoxyuridine injection

Adult fish were anaesthetized and placed on a wet sponge under a dissecting microscope. 5-bromo-2’-deoxyuridine (BrdU) was diluted in phosphate-buffered saline (PBS) to 2.5 mg/ml, and 30 µl were injected intraperitoneally 24 hours before heart dissection.

### Histology

Acid Fuchsin Orange G (AFOG) staining, *in situ* hybridization (ISH), fluorescent ISH, and whole-mount *in situ* hybridization (WM-ISH) were performed as described (Grivas et al., 2020; Münch et al., 2017). Primers for riboprobes used in this study are listed in Supplementary Table 2. For immunofluorescence, sections of paraffin-embedded tissue or cryosections were permeabilized with PBT (PBS with 0.01% TritonX-100) and washed with PBS before incubation with blocking solution (2% bovine serum albumin, 10% goat serum, and 2 mM MgCl_2_ in PBS). Sections were then incubated overnight at 4°’C with primary antibodies targeting BrdU (BD Transduction Laboratories), GFP (Aves Labs, Tigard), Aldh1a2 (Gene Tex), MEF2 (Santa Cruz Biotechnology, Santa Cruz, CA), phosho-Smad3 (Abcam, Cambridge, MA) or Mdka (Calinescu et al., 2009). Sections were then incubated with the appropriate secondary antibody and mounted after DAPI staining.

### Western Blot (WB)

Protein expression analysis by WB was performed as described (Grivas et al., 2020), using antibodies Mdka (Calinescu et al., 2009) and Alpha-Tubulin (Thermofisher scientific).

### Image analysis and quantification

To analyse cardiomyocyte proliferation, all MEF2^+^ and MEF2^+^/BrdU^+^ nuclei were counted in a 100 µm area around the injury. For the analysis of epicardial cell proliferation, all epicardial cells and epicardial/BrdU^+^ cells were counted. For the coronary endothelial cell proliferation analysis, GFP^+^ and GFP^+^/BrdU^+^ cells were counted within a 200 µm radius of the injury site. For psmad3 quantification, psmad3^+^ and psmad3^+^/epicardial cells were measured. The percentage was expressed as the MEF2^+^:MEF2^+^/BrdU^+^ ratio for cardiomyocytes, as the GFP^+^:GFP^+^/BrdU^+^ ratio for epicardial and endothelial cells, and psmad3^+^:psmad3^+^/epicardial cells for psmad3 quantification. To quantify regeneration, the injured (fibrotic tissue and collagen) and healthy (myocyte) areas were measured and expressed as a percentage of the total ventricular area. The percentage collagen:injury index was calculated as the ratio of the collagen area to the whole injury area. At least three sections each heart were used for all heart quantifications. The fin regeneration analysis, we measured the outgrowth of the regenerating fin from the amputation plane to the distal tip of the rays. All analyses were performed using Fiji (ImageJ, NIH).

### Gene expression analysis

Gene expression was analysed by qPCR using the power SYBR Green Master Mix (Applied Biosystems, Foster City, CA) and the ABI PRISM 7900HT Real-Time PCR System. Each condition was analysed using 3-4 biological replicates with 3 technical replicates per sample. Measurements were normalized to the expression of *18s* (in embryos) or of *elf1a* (in adult heart). All primers used in qPCR analysis are listed in Supplementary Table 3. RNA-seq analysis was conducted with 3 pools of 3 apexes from *mdka*^*+/+*^ or *mdka*^*cn105*^ hearts. cDNA libraries were prepared with the NEBNext Ultra II Directional RNA Library Prep Kit (New England Biolabs) and sequenced on a HiSeq 4000 system (Illumina) to generate 60 base single reads, and data were processed with RTA v1.18.66.3. FastQ files for each sample were obtained using bcl2fastq v2.20.0.422 software (Illumina). The resulting reads were mapped against the reference transcriptome GRCz11.99, and gene expression levels were estimated with RSEM (Li and Dewey, 2011). A single pairwise contrast was performed (KO vs WT). To increase the number of potentially relevant genes, *p-values* were not corrected for multiple testing, and changes in gene expression were considered significant if associated with a raw *P* value <0.05. qPCR provided complementary evidence for significant differential expression of selected genes, as described above. The resulting collection of 2029 differentially expressed genes was used for functional enrichment analysis with IPA to derive overrepresented gene lists from Ingenuity’s proprietary knowledge-base (IPAKB). IPAKB-derived gene lists consist of collections of genes belonging to the same signalling or metabolic pathway (Canonical Pathway analyses) that either are regulated by the same molecule (Upstream Regulator analyses) or are associated with the same disease or biological function (Downstream Effect analysis). Enrichments associated with a Benjamini-Hochberg adjusted *P* value <0.05 were considered significant. Depending on the number or the type of genes involved, IPA also issued predictions about the activation state of pathways or regulators in the form of a parameter called the z-score; activation or inhibition is indicated by positive or negative values, respectively. Circular plots, summarizing the relationship between selected regulators and target differentially expressed genes were generated with the R package GOplot (Walter et al., 2015).

### Accession number

Data are deposited in the NCBI GEO database under accession number ------. The following secure token has been created to allow review of record ------ while it maintains private status:

### Statistical analysis

Sample sizes, statistical tests, and *P* values are specified in the figure legends and were determined with GraphPad Prism software (GraphPad Software Inc., San Diego, CA). Statistical t tests were two-tailed. Differences were considered statistically significant at *P* <0.05.

## Acknowledgements

We thank E. Díaz at the CNIC animal facility for fish husbandry; B. Rios, V. García, and L. Méndez for technical support; the CNIC Microscopy Unit; P. Hitchcock for sharing the Mdka antibody; and S. Bartlett (CNIC) for English editing. RNA-seq was performed by the CNIC Genomics Unit. Data analysis was carried out by the CNIC Bioinformatics Units, and we want to specially thank Manuel J. Gómez for his help.

This work was supported by grants PID2019-104776RB-I00, CB16/11/00399 (CIBER CV) and RD16/0011/0021 (TERCEL) from the Spanish Ministry of Science, Innovation and Universities (MCIU) and grants from the Fundación BBVA (Ref.: BIO14_298), Fundación La Marató (Ref.: 20153431) and CardioNeT (Ref.: 28600) from the European Commission to J.L.d.l.P. D.G. held a PhD fellowship linked to grant CardioNeT. The cost of this publication was supported in part with funds from the European Regional Development Fund. The CNIC is supported by the Instituto de Salud Carlos III (ISCIII), the MCIU and the Pro CNIC Foundation, and is a Severo Ochoa Centre of Excellence (SEV-2015-0505).

## Author contributions

D.G. and A.G.-R. performed experiments. D.G. and J.L.d.l.P. designed experiments, reviewed all the data, prepared figures, and wrote the manuscript. All authors reviewed the manuscript during its preparation.

## Supplementary Figures and Legends

**Figure 1_Supplementary Figure 1.**
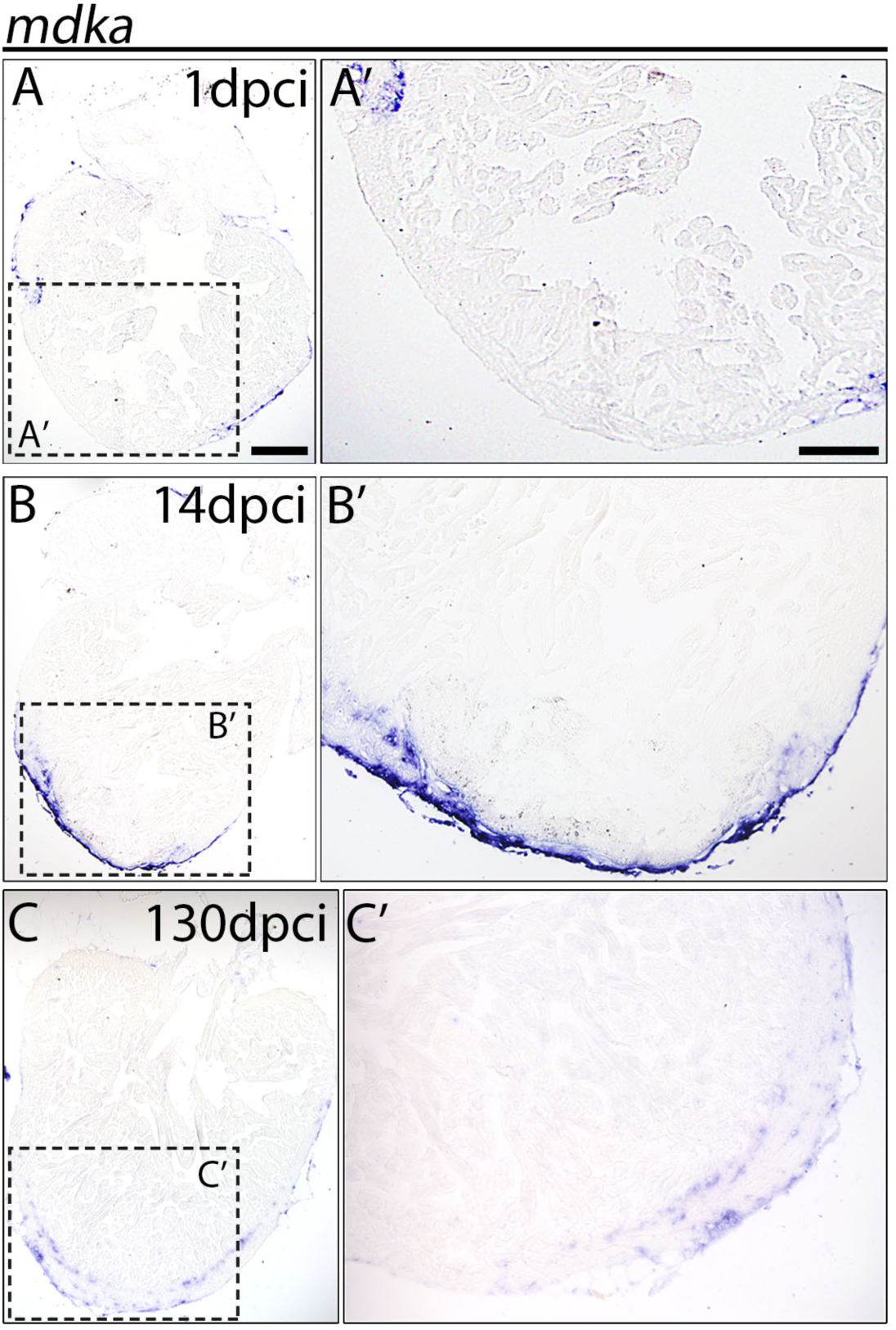
Expression of *mdka* in regenerating hearts. ISH of *mdka* in 1 dpci (A-A’), 14 dpci (B-B’), and 130 dpci (C-C’) hearts. Scale bars, 100 µm.

**Figure 2_Supplementary Figure 1.**
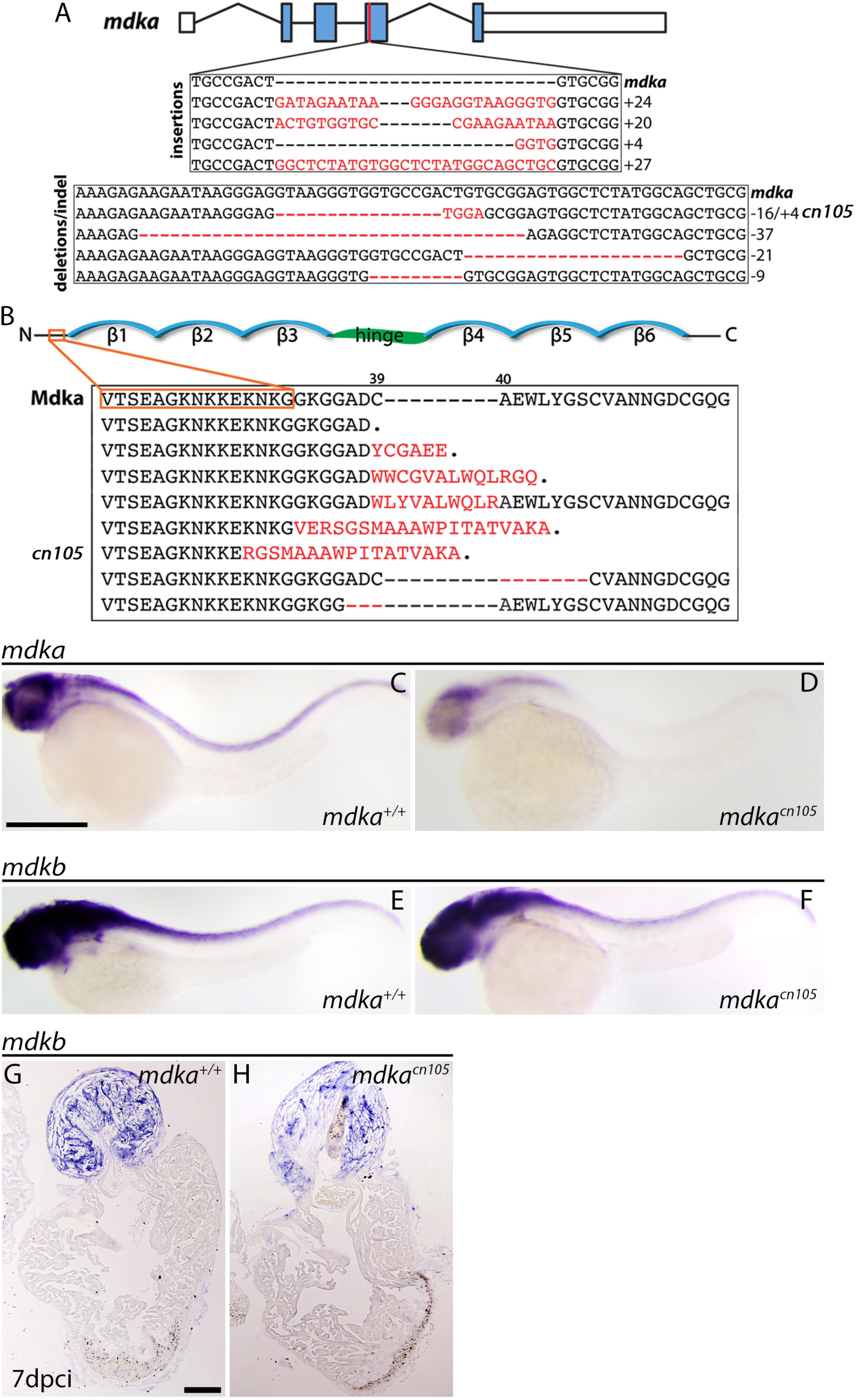
Analysis of *mdka* and *mdkb* in *mdka*^*cn105*^ embryos and injured hearts. (A) The *mdka* locus and DNA mutations introduced by CRISPR/Cas9. The red line indicates the CRISPER target site, red letters indicate insertions, and red hyphens deletions. (B) Mdka domain organization and predicted protein mutations. Red letters denote novel amino acids, red hyphen deletions. β stands for beta-strands and the hinge domain is shown in green. (C-D) Whole-mount ISH (WM-ISH) for *mdka* in 2-day post fertilization (dpf) *mdka*^*+/+*^ and *mdka*^*cn105*^ embryos. Scale bar, 200 µm. (E-F) WM-ISH of *mdkb* in 2 dpf *mdka*^*+/+*^ or *mdka*^*cn105*^ embryos. (G-H) ISH of *mdkb* in 7 dpci *mdka*^*+/+*^ or *mdka*^*cn105*^ hearts. Scale bar, 100 µm.

**Figure 3_Supplementary Figure 1.**
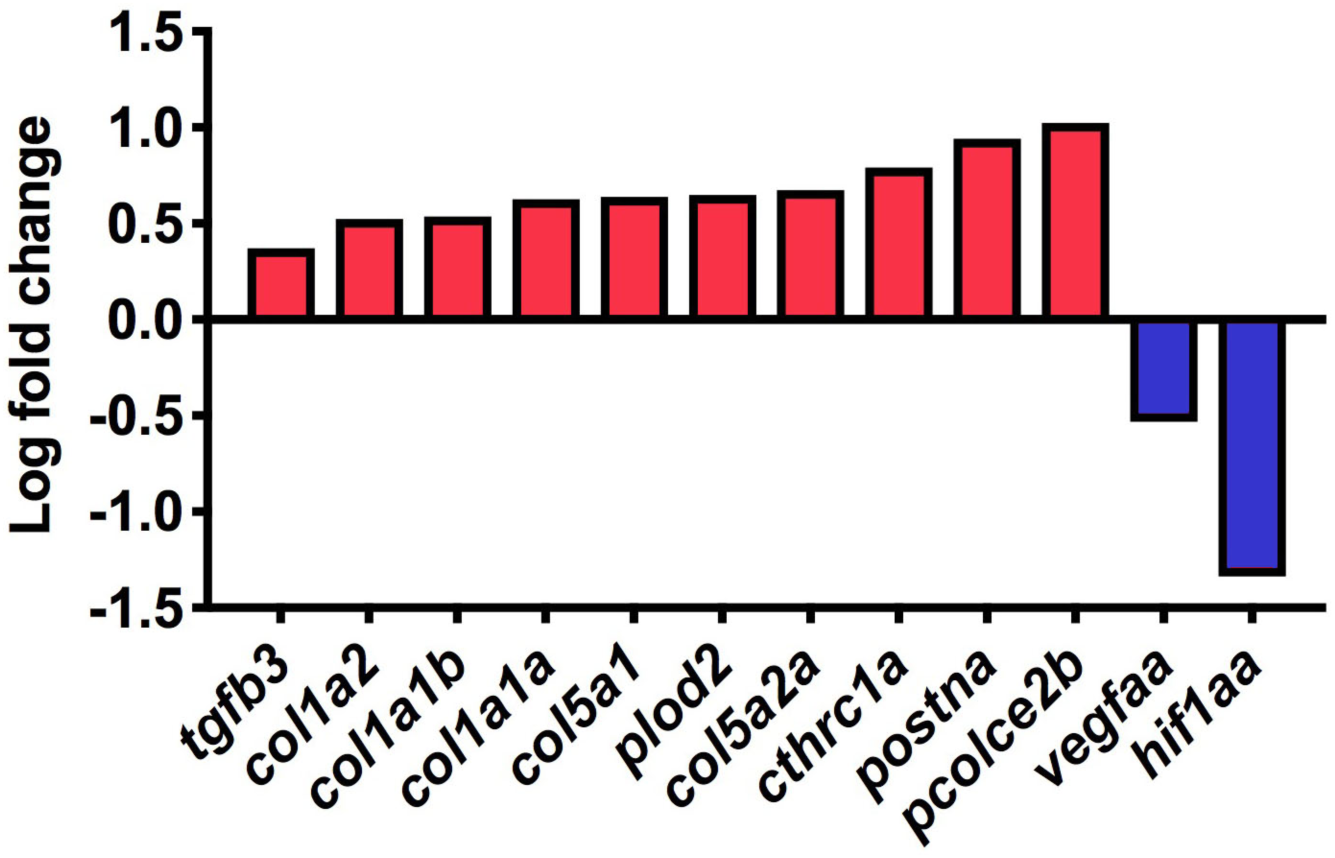
RNA-seq analysis of 7 dpci hearts. RNA-seq log fold changes of ECM components and genes important for angiogenesis.

**Figure 3_Supplementary Figure 2.**
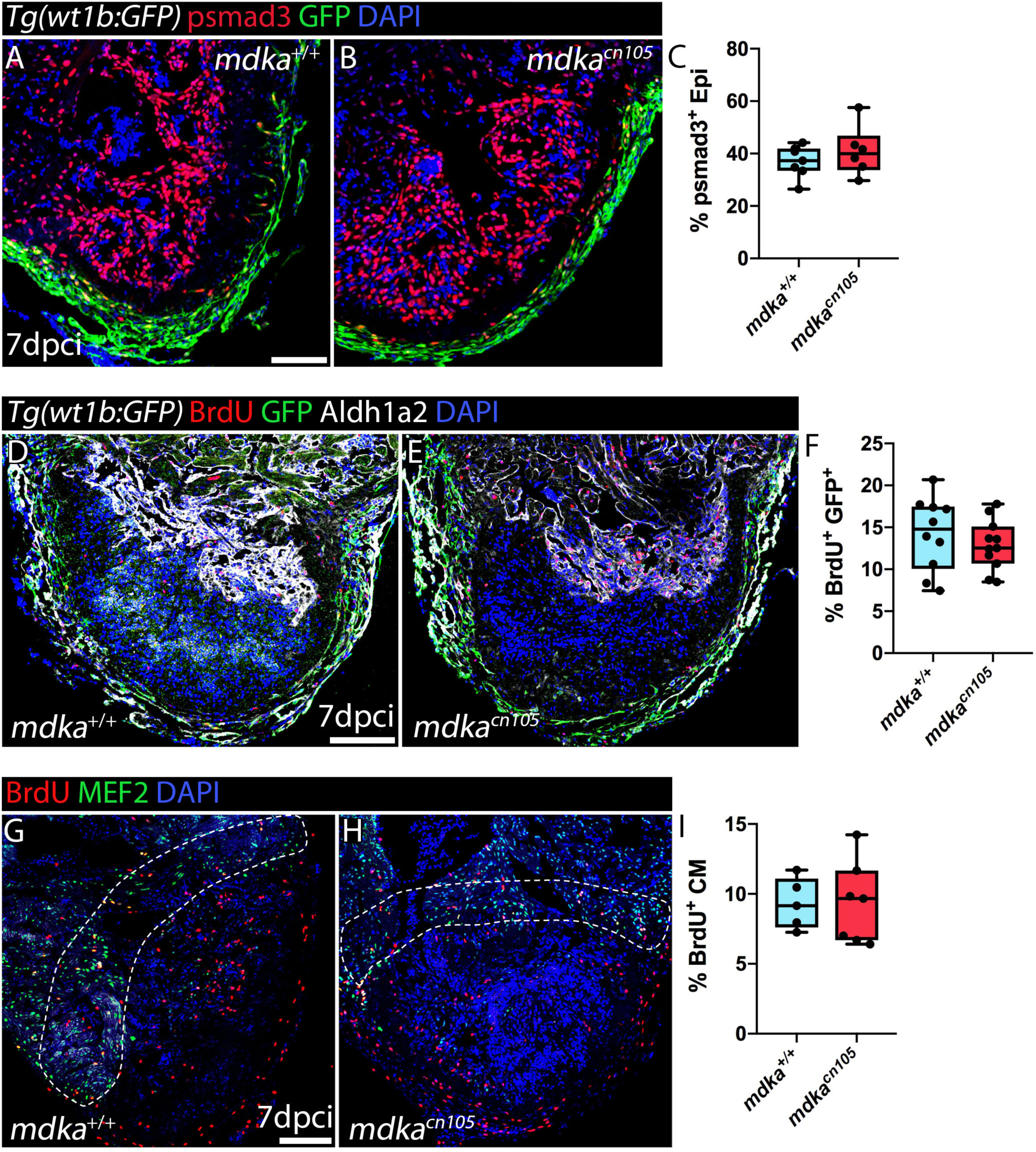
Epicardial cell and cardiomyocyte proliferation. (A, B) Immunofluorescence staining of *mdka*^*+/+*^ and *mdka*^*cn105*^ 7 dpci *Tg(wt1b:GFP)* heart sections for phospho-smad3 (psmad3, red) and GFP (green). (C) Quantification of psmad3^+^ epicardial cells. n_WT_ = 7, n_KO_= 6; t-test; Mean±S.D. (D, E) Immunofluorescence staining of *mdka*^*+/+*^ and *mdka*^*cn105*^ 7 dpci *Tg(wt1b:GFP)* heart sections for BrdU (red), GFP (green), and Aldh1a2 (white). (F) Quantification of BrdU^+^ epicardial cells. n_WT_ = 10, n_KO_= 11; t-test; Mean±S.D. (G, H) Immunofluorescence staining of BrdU and MEF2 in 7 dpci *mdka*^*+/+*^ and *mdka*^*cn105*^ hearts. (I) Quantification of BrdU^+^/MEF2^+^ cardiomyocytes. n_WT_ = 6, n_KO_= 7; t-test; Mean±S.D; Scale bars, 100 µm.

**Supplementary Table 2.**
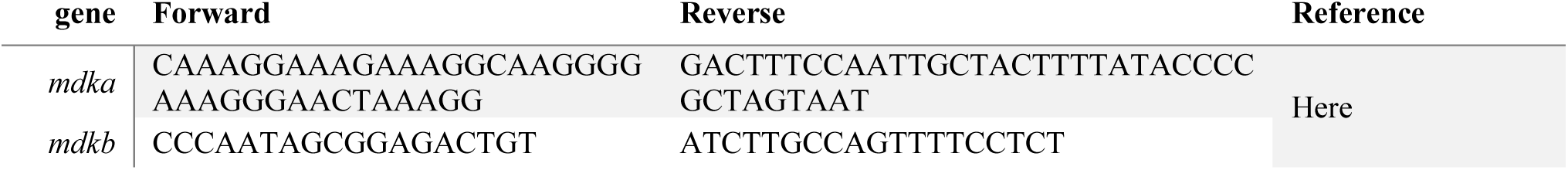
Primers for riboprobe.

**Supplementary Table 3.**
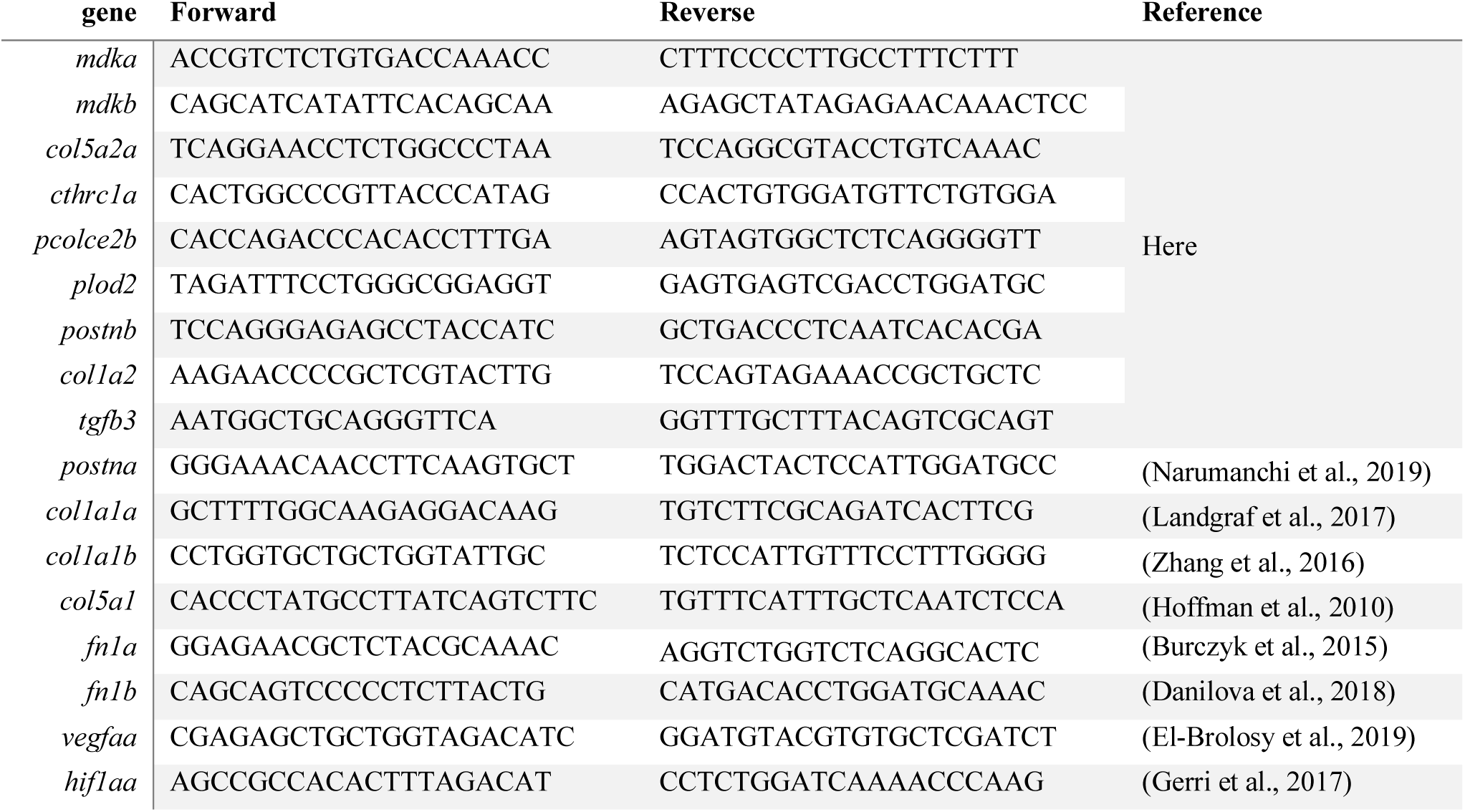
qPCR primers.

